# PLAGUE: Improved confidence plasmid reconstruction from short read assemblies

**DOI:** 10.64898/2026.07.10.737741

**Authors:** Karolina Kozubal, Faruk Dube, Amar Mujkic, Enrique Joffré, Lionel Guy, Helen Wang, Luisa W. Hugerth

## Abstract

As the vast majority of archival data is short-read based, reliable strategies for detecting and classifying plasmids out of these data are still needed. A widely adopted solution for binning plasmids with rich contextual data is MOB-Suite. However, a large portion of the available data has low coverage and results in fragmented assemblies. Under these circumstances, MOB-Suite delivers a high number of false positives, leading to wasteful follow-up experiments. To improve the reliability of MOB-Suite, while retaining and enriching its biological contextual output, we have developed PLAGUE. PLAGUE is a fast and low-memory consumption post-processing tool for MOB-Suite outputs, relying on checks of circularity, reads spanning the gap between the 3’ and 5’ ends and total coverage of the plasmid candidate. PLAGUE can reduce the number of false positives in MOB-Suite outputs by over 30%, while retaining almost full sensitivity. It is fully containerized and easy to use in clusters as well as locally.

## Introduction

Plasmids are mosaic mobile genetic elements. Rather than evolving as fixed, monolithic units, they are continuously assembled and reassembled from functional modules that move between plasmids and host chromosomes through duplication, recombination, transposition, and horizontal transfer. These functional modules (replicons, mobility and transfer machinery, and accessory cargo genes) form a dynamic architecture that makes plasmid reconstruction from sequencing data analogous to solving a puzzle whose pieces are constantly changing shape [1]. The problem is especially pronounced for small plasmids, which are frequently lost during genome assembly, and consequently remain the elements most likely to be misreconstructed or overlooked [2].

Accurate plasmid reconstruction is critical because plasmids are major vectors of antimicrobial resistance (AMR). By facilitating the horizontal transfer of antibiotic resistance genes (ARGs) across species and lineages, plasmids drive the dissemination of AMR at a pace that far exceeds that achievable through vertical chromosomal inheritance alone [3]. As a result, they play a central role in the emergence and spread of resistant pathogens, contributing directly to treatment failure and the global burden of AMR-associated mortality [4]. Consequently, understanding microbial evolution, resistance mechanisms, and epidemiological trends depends on the reliable identification and reconstruction of plasmids. Despite the growing adoption of long-read sequencing technologies, short-read sequencing remains the dominant source of openly available bacterial genomic data. As of 16th of June, 2026, the NCBI Sequence Read Archive held approximately 4.0 million short-read bacterial sequencing runs, with Illumina alone accounting for c. 3.95 million, compared with only <130,000 long-read runs from Oxford Nanopore and PacBio combined (**Table S1**). Long-read platforms thus represent roughly 3% of archival bacterial sequencing data, and short-read datasets outnumber long-reads by more than 30-fold. Robust and reliable bioinformatic approaches for plasmid reconstruction from short-read-based assemblies remain therefore a critical need.

Reference-based approaches are the predominant and highly effective strategy for plasmid identification from short-read assemblies, comparing contigs against curated databases to recognise known gene families such as replicons, relaxases, mobility genes, repetitive elements, or homologous plasmid sequences [5]. Their power derives from the substantial conservation of plasmid backbones, which makes catalogued references broadly informative. Recombination, however, means that the plasmids circulating in nature are partly moving targets: backbones are shuffled and reassembled over time, so a query element may diverge from its closest reference even as new recombinants arise. This does not undermine the reference-based strategy so much as define where its predictions carry more uncertainty, typically at the margins, where partial reconstruction or chromosomal contamination inflate the false-positive rate. Paganini et al. report 40.1% of chromosomal contamination after using MOB-suite, a plasmid reconstruction tool from WGS assemblies [6]. Such cases are well suited to downstream validation rather than a change of strategy, motivating a refinement layer that retains the strengths of reference-based reconstruction while resolving its less confident calls.

Compounding this, bacteria typically carry several plasmids simultaneously, so a single assembly must be resolved into multiple distinct elements. MOB-suite [5] is a widely used reference-based tool built for exactly this problem. It integrates MOB-recon for plasmid detection, MOB-typer for typing, and MOB-cluster for clustering, and it works in two stages: it first classifies contigs against well-curated reference databases by their backbone signatures, then reconstructs and bins them into candidate plasmids. This two-step design distinguishes it from pure classifiers such as geNomad [7], which assign contigs a plasmid-or-not label but do not attempt per-plasmid reconstruction.The binning that MOB-suite provides, recording which contigs belong to which reconstructed plasmid, is precisely what makes its output valuable for downstream evolutionary and epidemiological analysis.

MOB-suite’s reported performance is strong, with MOB-recon described in the original publication as demonstrating both high sensitivity and specificity, at 95 % and 88 % respectively. In practice, however, its predictions frequently include false positives, particularly when applied without additional validation. In our own analyses of Illumina short-read datasets retrieved from NCBI, a substantial fraction of predicted “plasmids” were very short contigs that did not resemble plausible plasmids and were better explained as assembly artefacts. This gap between catalogue performance and real-world output points to a need for more precise and comprehensive validation downstream of MOB-suite.

The ideal plasmid reconstruction workflow would retain the rich annotation and per-plasmid binning provided by tools such as MOB-suite while sharply reducing its false-positive rate. To this end, we developed PLAGUE (<u>PL</u>asmid <u>A</u>ssembly <u>G</u>enetic <u>U</u>nit <u>E</u>valuator), a post-processing pipeline that extendsMOB-suite with a three-step validation framework: (i) circularisation detection using Circlator’s minimus2 module [8], (ii) analysis of discordant read pairs at contig termini, and (iii) coverage assessment from BAM files. Together, these complementary metrics provide evidence for coverage-based filtering and structural confidence. PLAGUE is designed for short-read data spanning a broad range of sequencing depths and qualities, including surveillance and outbreak investigations, datasets at moderate-to-low coverage, and routine Illumina sequencing projects. Importantly, the framework remains conservative when applied to high-quality assemblies, preserving well-supported predictions while improving confidence in low-quality datasets.

Because PLAGUE inherits and preserves MOB-suite’s per-plasmid binning information, each output record retains the contig-to-plasmid mapping required for downstream evolutionary and epidemiological analyses. The resulting output remains directly compatible with plasmid-network tools such as PLING [9], enabling reconstructed plasmids to be tracked across isolates rather than analysed as disconnected contigs. By combining circularisation verification, coverage assessment, and discordant-read detection, PLAGUE reduces false-positive predictions while producing an annotated results table that supports downstream biological and epidemiological analysis. We further benchmark PLAGUE against a geNomad-based post-filtering strategy, providing comparison with a widely adopted and methodologically distinct approach to plasmid classification.

## Methods

### Pipeline implementation

The PLAGUE pipeline was implemented in Nextflow for scalable, portable, and reproducible execution across computing environments [10]. It takes assembled bacterial genomes (FASTA) as primary input, with paired-end short-read FASTQ files used for read mapping and coverage analysis. All intermediate files and logs are retained to enable manual inspection and re-analysis. The workflow is organized into sequential modules producing standardized intermediate outputs. MOB-suite identifies putative plasmid contigs from the input assemblies and reports plasmid annotations, such as replicon types and mobility metadata. ABRicate [11] is run against the NCBI AMRFinderPlus database [12] (default) to annotate resistance determinants on predicted plasmid contigs. SeqKit2 [13] computes contig-level statistics and flagged single-contig plasmids for downstream structural validation. All data is compiled and reported in a structured table to enable manual assessment and decisions where needed (**Supplementary Table S2**).

#### Filters applied to single-contig plasmids

The bulk of the validation is aimed at single-contig plasmids. Single-contig predictions are the most error-prone class of MOB-suite output: a short, single contig carries few database-detectable elements, so a misclassified chromosomal fragment is hard to distinguish from a genuine small plasmid. Multi-contig reconstructions, by contrast, aggregate more sequence and therefore more corroborating evidence (replicon, relaxase, and mobility signals across several contigs), making them far less prone to misclassification. The bulk of PLAGUE’s validation is therefore directed at single-contig plasmids.

Circlator [8] (using the minimus2 module) assesses contig circularity by splitting each contig into two halves and testing whether the 5′ and 3′ ends share an overlapping sequence. If an overlap is detected, Minimus2 merges the halves to produce a single circularized sequence. Circlator outputs, including circularization status, are retained for every contig. Next, short reads are mapped to plasmid-derived contigs using BBMap [14], and per-base coverage is summarized with samtools depth [15] to derive mean coverage; sorted BAM files are retained for downstream analyses (Fig. 1).

**Figure 1:**
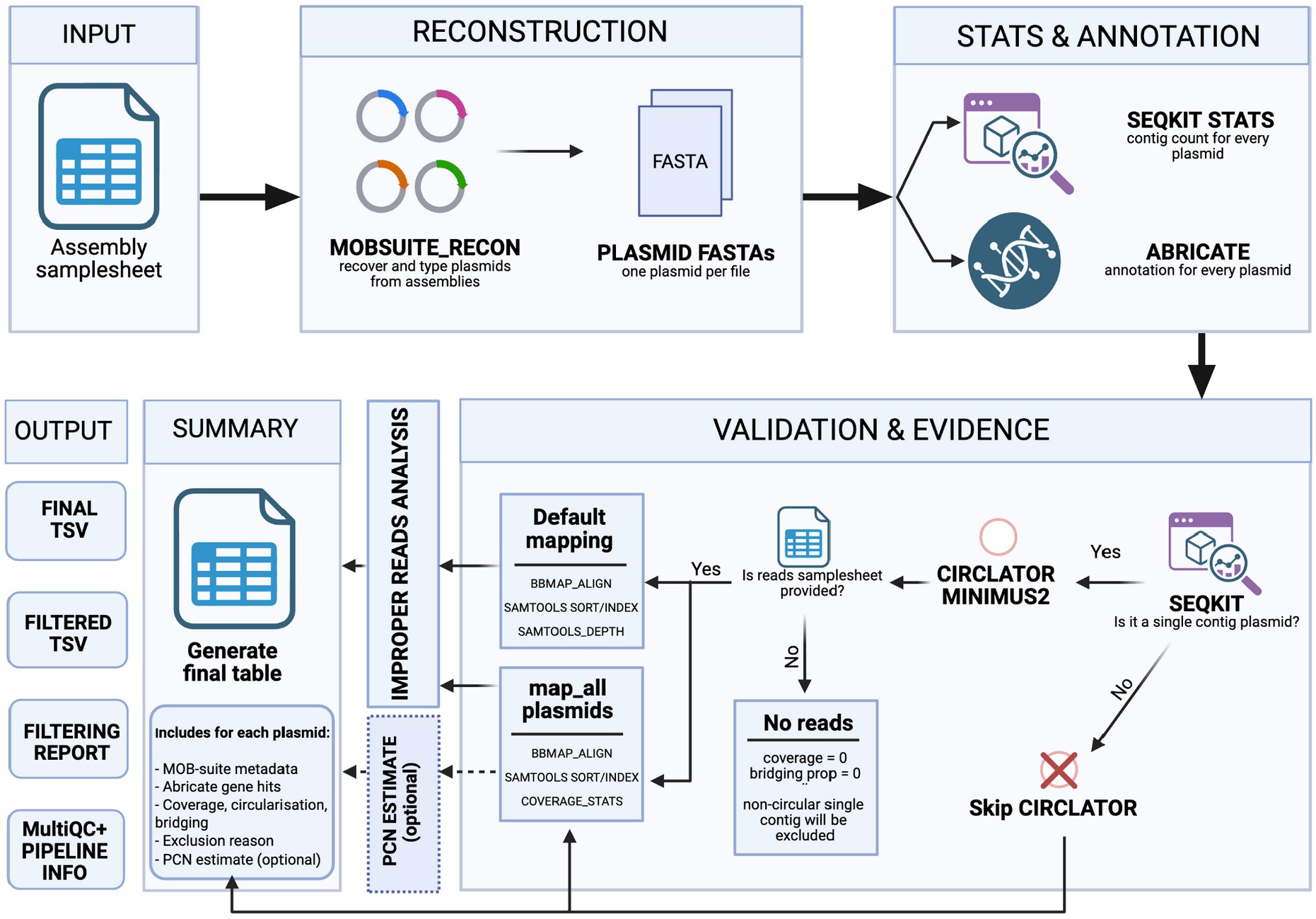
Summary of the PLAGUE pipeline and its outputs.

Because contigs may be circular without being fully assembled, an additional circularity analysis is performed, as follows. discordant read pairs near contig ends (300 bp by default) are identified from alignments using SAMtools (v1.22.1) by detecting reads that are mapped, with a mapped mate, but not properly paired. These improper reads, when present in both 5’ and 3’ terminus windows, are classified as terminus-bridging reads. The proportion of improper read pairs and the number of terminus-bridging reads per contig serve as complementary metrics to flag potential misassemblies and to corroborate circularity calls made by Circlator (Fig. 1).

Each single-contig plasmid was evaluated against three complementary filters (Filters A–C) that combine circularity, coverage depth, and improper-read proportion (Fig. 1). The thresholds were chosen empirically from the benchmark data (detailed below). Filter A removes contigs that lack circular support and have low coverage - a negative Circlator circularity call combined with a mean coverage below 25x. Filter B removes contigs with very low coverage and high read discordance at their termini - a mean coverage below 10x combined with an improper-read proportion above 98%. Filter C removes contigs that lack circularity evidence yet show minimal discordant-read signal - a negative Circlator call combined with an improper-read proportion below 10%, indicating no detectable junction.

Each single-contig plasmid is tested against all three filters. Three outputs are produced: plague_final_results.tsv (all predictions, with metrics, annotations and a per-plasmid exclusion_reason), plague_filtered_results.tsv (the high-confidence set: only plasmids passing every filter), and plague_filtering_report.txt (aggregate counts: total, validated and excluded plasmids, plus per-filter exclusion counts). SeqKit2 stats --all is computed per plasmid but used only to assign contig numbers (single-versus multi-contig plasmid). The output table consolidates, per plasmid, the MOB-typer annotations (predicted mobility, replicon and relaxase type(s), MPF type, oriT type, MASH nearest neighbour and predicted host-range taxon) together with Abricate AMR genes, coverage, circularisation status, bridging-read proportion and, when enabled, plasmid copy number. Multi-contig plasmids, which cannot be circularity-validated, are never excluded, but are presented to the user with full contextual information including per-contig coverage, empowering users to make their own interpretation of these plasmids’ accuracy. An example output table is presented in supplementary **Supplementary Table S2**.

#### Determining default settings

To determine default settings, each candidate signal (contig length, mean coverage, and the proportion of improperly paired reads), was treated as a single-feature classifier and its cut-off was swept across the observed range. Sensitivity and F1 were computed against the RefSeq-containment ground truth, selecting for each signal the value that maximised F1 while preserving sensitivity. For the initial filtering threshold, 30 kb in length was the point at which full-filter F1 peaked. For coverage, the 25x cut sits at the top of an F1 plateau spanning 25–30x while retaining higher sensitivity than the 30x alternative (0.974 versus 0.949). On the other hand, a mean coverage below 10x combined with an improper-read proportion above 98% triggers plasmid removal as a safeguard against grossly misassembled contigs, despite not having removed any plasmid candidates in the present benchmark. Finally, for circularity, a negative Circlator call combined with an improper-read proportion below 10%, indicates no detectable junction, with the 10% threshold taken as the conservative lower end of an F1 plateau spanning 5–25%. The full selective parameter sweep is presented in **Supplementary Table S3**.

### Validation

#### Validation dataset and preprocessing

A total of 41 *Escherichia coli*, 36 *Staphylococcus aureus*, and 25 *Bacillus cereus* Illumina-sequenced genomes were retrieved from the NCBI RefSeq database using NCBI Datasets [16] (**Table S4-S6**). All isolates were processed with BACTpipe v3.1.0 using default settings (Rudbeck et al., 2021). Raw paired-end reads were preprocessed with fastp [17] for quality control, adaptor trimming, and deduplication; taxonomic classification and contamination screening were performed with Kraken2 [18]; and genome assemblies were generated with Shovill [19] (which internally invokes SPAdes [20] to produce contig FASTA files. The resulting assemblies and their corresponding raw reads were used as input to the PLAGUE pipeline.

#### Coverage downsampling

To evaluate pipeline robustness across varying sequencing depths, the original short-read datasets were subjected to controlled downsampling using reformat.sh from the BBMap suite [14]. Four coverage tiers were generated for each isolate: the original coverage, 30x, 20x, and 10x. This design captures a spectrum of sequencing depths from high-coverage benchmarks to the lower limits of assembly viability, reflecting the suboptimal depth frequently encountered in publicly archived datasets, where reduced coverage often produces highly fragmented assemblies that confound traditional plasmid prediction tools. For each coverage tier, every MOB-suite plasmid prediction was assigned a PLAGUE verdict (STAY or OUT) based on the filtering criteria described above.

#### Reference comparison using Mash

To enable high-throughput comparison between pipeline predictions and the NCBI RefSeq [21] ground truth, a k-mer-based approach was implemented using Mash [22] v2.3. For each isolate, the reference plasmid sequences were subset from the complete RefSeq assembly using SeqKit2 split [13], producing isolate-specific reference sketches. An analogous set of isolate-specific sketches was constructed from the MOB-suite reconstructed plasmids. Containment metrics were then calculated using mash screen, yielding containment on the reference (CR = |B∩R|/|R|) and containment on the assembly (CA = |B∩R|/|B|), where B is a MOB-suite bin and R is an NCBI reference plasmid. To derive a single summary value per MOB-suite bin, the ratio CR/CA = |B|/|R| was computed, capturing the relative size of the predicted bin with respect to the reference. The ratio thresholds were: tau = 0.50, 0.80, 0.95. These values were integrated into a master data frame alongside MOB-suite and geNomad predictions, run with geNomad end-to-end command with default options [7].

Because geNomad operates as a per-contig classifier rather than a binning tool, contig names were extracted from each MOB-suite bin and matched to the corresponding geNomad predictions. Since Mash screen was applied at the bin level, multi-contig plasmids received a single CR and CA value. To enable per-plasmid comparison at the bin level, geNomad contig-level plasmid scores were aggregated using a length-weighted mean: lw = Σ(score_i × length_i) / Σ length_i, where score_i and length_i denote the plasmid score and sequence length of the i-th contig within the bin.

#### Performance evaluation

Each prediction from MOB-suite [5], geNomad [7], and the PLAGUE-refined pipeline was classified as a true positive (TP), false positive (FP), or false negative (FN). True negatives were excluded from the analysis, as they could only be drawn from a post-MOB-suite conditional distribution and would therefore not constitute an unbiased sample of the broader sequence space. A MOB-suite prediction was counted as a TP if computed ratio CR/CA = |B|/| R| was high enough (tau = 0.50, 0.80, 0.95), an FP if the ratio was smaller then the threshold ratio, and an FN if a known reference plasmid had no corresponding MOB-suite cluster. For geNomad, a contig or bin scoring above the plasmid classification threshold was counted as a TP if it matched a known NCBI accession, an FP if it did not, and an FN if a verified plasmid received a non-plasmid classification or was not detected. For the PLAGUE-refined pipeline, a STAY verdict matching a verified NCBI accession was counted as a TP, a STAY verdict lacking an NCBI match as an FP, and a verified plasmid incorrectly assigned a non-STAY verdict or missed entirely as an FN. These operational parameters are summarized in Table 1.

**Table 1.**
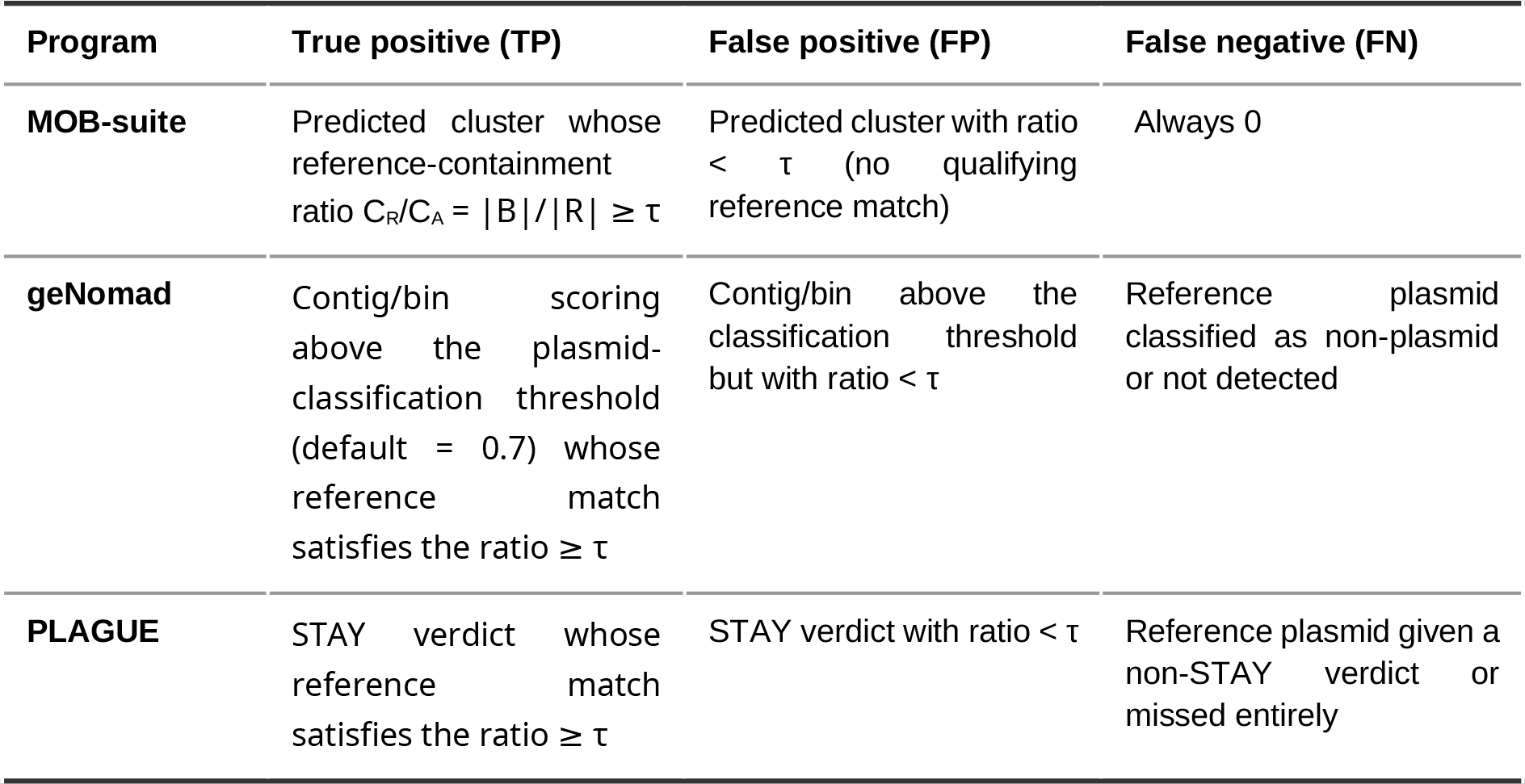
Classification of predictions as true positives (TP), false positives (FP), and false negatives (FN) for each program, relative to the NCBI RefSeq ground truth. A prediction is counted as a true match when the Mash containment ratio C_R_/C_A_ = |B|/|R| reaches the threshold τ, evaluated at τ = 0.50, 0.80, and 0.95; B denotes a predicted bin and R the matched RefSeq reference plasmid. The same τ threshold defines a true match for all three programs. geNomad contig-level scores were aggregated to bin level by a length-weighted mean before thresholding. True negatives were excluded, as they could only be drawn from a post-MOB-suite conditional distribution. Because PLAGUE operates only on MOB-suite bins, its true positives are a subset of MOB-suite’s, making MOB-suite the upper bound on sensitivity.

Precision, sensitivity, and F1 score were calculated across all four coverage tiers as follows: Sensitivity = TP / (TP + FN), Precision = TP / (TP + FP), and F1 = 2 · Sensitivity · Precision / (Sensitivity + Precision). These metrics allow direct quantification of the degree to which the PLAGUE filtering layer improves precision by reducing false positives in fragmented, low-coverage assemblies.

## Results

### Overview: PLAGUE filters false positives while preserving sensitivity

To evaluate PLAGUE, the pipeline was benchmarked against raw MOB-suite output and against geNomad, an independent, machine-learning-based plasmid classifier, across three bacterial species (*Escherichia coli, Staphylococcus aureus*, and *Bacillus cereus*) at four sequencing depth tiers (10x, 20x, 30x, and original coverage; hereafter OG). Performance was measured by sensitivity (true-positive rate), precision (positive predictive value, PPV), and F1 score relative to NCBI RefSeq plasmid annotations used as ground truth. Because PLAGUE operates as a post-processing layer on top of MOB-suite, MOB-suite has sensitivity 1.000 by construction: every true positive considered in this benchmark originates from MOB-suite output, making it the starting point and upper bound for recall rather than a competing method. The key question is therefore how much of this baseline sensitivity PLAGUE preserves after filtering, and at what gain in precision. Across all tiers and species, unfiltered MOB-suite output produced large numbers of false positives, with PPV values as low as 0.120 and F1 scores as low as 0.214. PLAGUE’s filtering layer consistently reduced this false-positive burden while retaining the large majority of genuine plasmid predictions, demonstrating that a substantially cleaner plasmid set can be obtained at modest and predictable sensitivity cost.

### *Escherichia coli:* high precision gain at minimal sensitivity cost

On the *E. coli* dataset, comprising 41 RefSeq-validated isolates (**Supplementary Table S4**), at original sequencing depth, the MOB-suite baseline comprised 179 true positives and 41 false positives (PPV = 0.814, F1 = 0.897). PLAGUE filtering reduced false positives from 41 to 36 while removing only 7 genuine predictions, yielding Sensitivity = 0.961, PPV = 0.827, and F1 = 0.889 (**Fig. 2A-B**). PLAGUE therefore retained 96.1% of all true positives from the MOB-suite baseline while eliminating 12% of its false positives: a conservative, targeted improvement that preserves nearly the complete true-positive signal.

**Figure 2.**
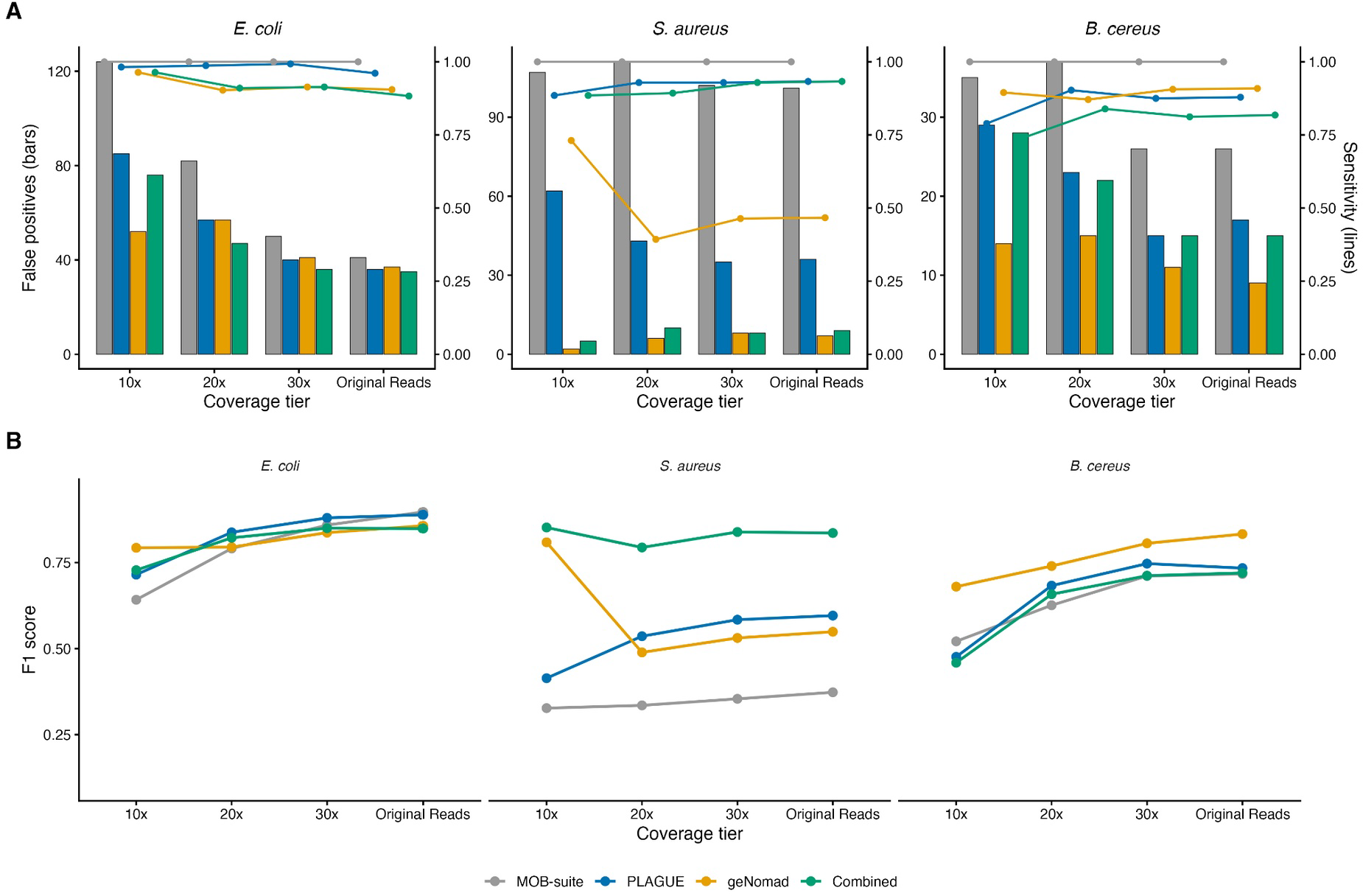
Benchmarking plasmid identification performance across coverage tiers and bacterial hosts. **(A)** False positive counts (left y-axis, grouped bars) and sensitivity percentages (right y-axis, trend lines) broken down by target species (*Escherichia coli, Staphylococcus aureus*, and *Bacillus cereus)* across sequencing depths (10x, 20x, 30x) and Original Reads. Bars and point elements are shaded by tool category (MOB-suite: grey, PLAGUE: blue, geNomad: yellow, Combined: green). **(B)** Overall F1 score trajectories across coverage tiers for target species

Performance gains from PLAGUE were pronounced and consistent across all downsampled tiers. At 30x coverage, PLAGUE raised F1 from 0.859 to 0.880 and reduced false positives from 50 to 40, while retaining 151 of 152 true positives (Sensitivity = 0.993). At 20×, F1 improved from 0.791 to 0.838 (FP reduced from 82 to 57; Sensitivity = 0.987). The most striking absolute gains arose at 10x coverage, where assembly fragmentation dramatically inflated the MOB-suite false-positive set: 124 false positives against only 111 true positives (PPV = 0.472, F1 = 0.642). PLAGUE reduced false positives by 31% to 85, raising F1 to 0.715, while losing only 2 of 111 true positives (Sensitivity = 0.982). Aggregated across all four coverage tiers, PLAGUE reduced the cumulative false-positive count from 297 to 218, a 27% reduction, while retaining 585 of 597 true positives (Sensitivity = 0.980) and improving overall F1 from 0.801 to 0.836 (**Fig. 2A-B**).

Notably, for *E. coli*, PLAGUE alone achieved a higher F1 than geNomad alone across all tiers combined (0.836 vs 0.823), principally because PLAGUE retained substantially higher sensitivity (0.980 vs 0.918) despite a marginally lower precision (PPV 0.729 vs 0.746). The combined PLAGUE & geNomad mode yielded only marginal additional precision gains for this species, suggesting that PLAGUE’s structural and coverage-based filters are already well-calibrated for the fragmentation patterns typical of *E. coli* short-read assemblies.

### *Staphylococcus aureus*: high false-positive rate

The 36 *Staphylococcus aureus* assemblies (**Supplementary table S5**) presented the most challenging benchmarking scenario: MOB-suite predictions were dominated by false positives at every coverage tier, a pattern consistent with the high degree of assembly fragmentation and contig misclassification typical of this pathogen’s complex plasmid content. At original coverage, the MOB-suite baseline comprised 30 true positives and 101 false positives (PPV = 0.229, F1 = 0.373), rendering unfiltered output practically uninformative for downstream analyses. PLAGUE’s filtering reduced false positives to 36, eliminating two thirds of the FP count, and raised precision to 0.438 and F1 to 0.596, while retaining 28 of 30 true positives (Sensitivity = 0.933; **Fig 2 C-D**).

At the lowest depth tier (10x), where the MOB-suite baseline comprised 26 true positives and 107 false positives (PPV = 0.195, F1 = 0.327), PLAGUE reduced false positives to 62, a 42% reduction, while maintaining Sensitivity = 0.885. Summed across all tiers, PLAGUE reduced the total false-positive count from 421 to 176 (a 58% reduction) with a sensitivity of 0.920 (**Fig. 2C-D**). This combination of large FP reduction and near-intact sensitivity is the most compelling demonstration of PLAGUE’s value: even when the underlying assemblies are of poor quality, the pipeline eliminates the majority of spurious predictions without discarding genuine plasmids.

For *S. aureus*, the combined PLAGUE & geNomad mode provided the strongest overall performance. At original coverage, with a geNomad length-weighted threshold of 0.20, the combined mode achieved Sensitivity = 0.933, PPV = 0.757, and F1 = 0.836, compared to F1 = 0.596 for PLAGUE alone and F1 = 0.549 for geNomad alone. Aggregated across all tiers, the combined mode reached Sensitivity = 0.911 and F1 = 0.829 (**Fig. 2C-D**). This result indicates that for species with high intrinsic FP rates, the two complementary evidence streams, structural/coverage-based filtering from PLAGUE and sequence-based scoring from geNomad, are mutually reinforcing.

### *Bacillus cereus*: moderate precision improvement

*Bacillus cereus* (**supplementary table S6)** exhibited an intermediate false-positive rate relative to *S. aureus*. At original coverage, the MOB-suite baseline comprised 33 true positives and 26 false positives (PPV = 0.559, F1 = 0.717). PLAGUE reduced false positives to 17, improving precision to 0.630 and F1 to 0.734, while retaining 29 of 33 true positives (Sensitivity = 0.879; **Fig. 2E-F**). Gains were consistent across downsampled tiers: at 30x, PLAGUE raised F1 from 0.711 to 0.747 (FP reduced from 26 to 15, Sensitivity = 0.875), and at 20x, F1 improved from 0.626 to 0.683 (FP reduced from 37 to 23, Sensitivity = 0.903). Aggregated across all tiers, PLAGUE reduced total false positives from 124 to 84 - a 32% reduction - with an overall sensitivity of 0.870.

Notably, for B. cereus, geNomad alone outperformed PLAGUE alone across aggregated tiers (F1 = 0.772 vs 0.669), with geNomad achieving higher precision at the cost of only marginally lower sensitivity (0.896 vs 0.870 for PLAGUE). This contrast suggests that geNomad’s sequence-based classifier is particularly effective for Bacillus plasmids, possibly because their sequence features are sufficiently distinctive to allow reliable classification without structural evidence. However, the combined PLAGUE & geNomad mode did not outperform geNomad alone for this species (aggregated F1 = 0.646 vs 0.772), indicating that the PLAGUE filter added conservatism that removed some true positives that geNomad correctly retained. For B. cereus, users prioritising precision may therefore prefer to run geNomad as a standalone post-processor, or to apply PLAGUE with relaxed filter thresholds.

### Robustness across sequencing depths

A key design requirement for PLAGUE is that it should remain accurate across a wide range of sequencing depths, as publicly archived datasets frequently include samples at sub-optimal coverage. The benchmark results confirm that PLAGUE’s filtering logic scales appropriately with depth across all three species. At 10× coverage - the lowest tier examined, at which assembly fragmentation is most pronounced and MOB-suite false-positive rates peak - PLAGUE consistently reduced FPs relative to unfiltered output: by 31% in *E. coli* (124 to 85), by 42% in *S. aureus* (107 to 62), and by 17% in *B. cereus* (35 to 29). Sensitivity at 10x remained high in *E. coli* (0.982) and *S. aureus* (0.885), with the larger sensitivity cost in *B. cereus* (0.789) likely reflecting genuine borderline predictions that lose structural signal at this depth.

Equally important, PLAGUE imposes no meaningful penalty on high-quality, full-coverage assemblies. At original depth, sensitivity in *E. coli* is 0.961, recovering 172 of 179 baseline true positives: a loss of only 7 plasmids. In *S. aureus* and *B. cereus*, sensitivity at original depth is 0.933 and 0.879, respectively. The few predicted plasmids removed at full coverage are most likely genuine false positives retained by MOB-suite without PLAGUE’s structural validation, deriving from fragmented or low-coverage artefacts. This conservative behaviour at high depth is a deliberate consequence of PLAGUE’s multi-criterion filter design: a prediction is only removed when multiple independent lines of evidence (circularisation failure, coverage collapse, and discordant-read accumulation) converge on the same contig, making inadvertent removal of well-supported true positives unlikely.

### PLAGUE and geNomad operate as complementary tools

Comparing PLAGUE with geNomad across all species reveals a consistent pattern: PLAGUE prioritises sensitivity while geNomad prioritises precision, and the two tools are therefore complementary rather than interchangeable. For *E. coli*, PLAGUE outperformed geNomad in sensitivity (0.980 vs 0.918, aggregated) and in F1 (0.836 vs 0.823), suggesting that geNomad’s classifier may penalise novel or divergent *E. coli* plasmid sequences not well-represented in its training data. For *S. aureus*, geNomad alone achieved higher precision than PLAGUE alone (PPV 0.713 vs 0.369, aggregated), though at substantially reduced sensitivity (0.509 vs 0.920); the combined mode reconciled these trade-offs most effectively (Sensitivity = 0.911, PPV = 0.761, F1 = 0.829). In *B. cereus*, geNomad alone performed best overall (F1 = 0.772 vs 0.669 for PLAGUE), and the combination did not add value, likely because the structural filter discarded predictions that geNomad correctly retained at the sequence level. These two approaches also complement each other in terms of computational needs: while geNomad’s peak memory requirement is c. 6-fold that of PLAGUE, it is also up to 4 times faster in wall-clock time, with comparable CPU requirements **(Fig. 3)**.

**Figure 3.**
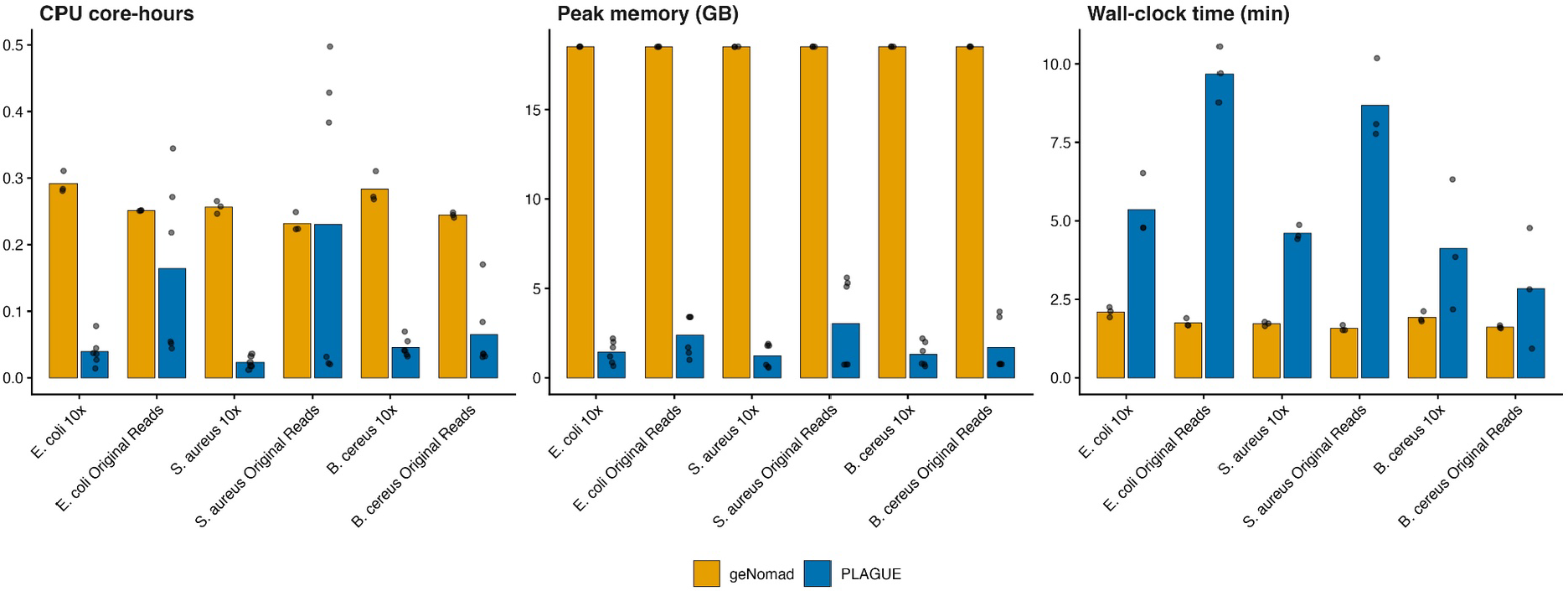
Computational resource profiles for plasmid identification across tools. Bars represent the mean metrics calculated for 10x coverage as well as original sequencing reads for *Escherichia coli, Staphylococcus aureus*, and *Bacillus cereus*; points represent individual sample replicates. Evaluation panels compare cumulative processor overhead (CPU core-hours), peak memory consumption (Peak memory, GB), and wall-clock processing duration (Wall-clock time, min). PLAGUE (blue) demonstrates a lightweight memory footprint at the expense of higher execution times, whereas geNomad (yellow) delivers rapid runtimes with elevated peak memory requirements.

These results indicate that PLAGUE is best positioned as a default post-processing step after MOB-suite, suited to users who wish to eliminate the most egregious false positives, particularly from low-coverage or fragmented assemblies, while preserving essentially the full complement of true plasmid predictions. The additional combined mode (PLAGUE & geNomad) offers a higher-precision alternative for applications where a clean, annotated plasmid set is more important than exhaustive recall, and is particularly effective for *S. aureus*. For species such as *B. cereus* where sequence-based evidence is independently informative, geNomad can complement or substitute PLAGUE’s structural filters, depending on the research context. In all scenarios, PLAGUE’s configurable filter thresholds allow users to tune the precision-sensitivity trade-off for their specific dataset characteristics. Even without automatic filtering, PLAGUE’s rich output table structure (**Supplementary Table S2**) can add important context to the assembly results.

## Discussion

Accurate plasmid identification and reconstruction is a cornerstone of genomic epidemiology as plasmids drive bacterial adaptation, facilitate the horizontal dissemination of antimicrobial resistance [3], and frequently carry virulence and fitness determinants that shape pathogen evolution. Reliable plasmid reconstruction therefore strengthens surveillance, outbreak investigations and research alike. Although long-read sequencing technologies continue to expand, short-read sequencing remains, by a substantial margin, the dominant source of publicly available bacterial genomes, with Illumina alone accounting for 95.4% of bacterial sequencing runs in the NCBI sequence Read Archive (Supplementary Table S1). Consequently, accurate reconstructions from these fragmented, short-read assemblies remains the prevailing challenge.

MOB-suite [5] is currently the most widely adopted reference-based tool for this task, valued for its integrated plasmid typing, clustering and per-plasmid binning capabilities. The original benchmark reported a 95% sensitivity and 88% specificity. However, performance often deteriorates when applied to fragmented short-reads assemblies; for example, a recent study reported 40.1% chromosomal contamination among MOB-suite-derived plasmid prediction in *E. coli* [6]. Our own benchmark reproduced this pattern, with unfiltered MOB-suite precision (PPV) falling as low as 12% and F1 as low as 21% in the hardest species-coverage combinations tested. The broader plasmid-detection landscape is active and methodologically diverse, spanning graph-based binners such as HyAsP [23], PlasmidEC/gplas2 [24] and PlasBin-flow [25], supervised classifiers such as mlplasmids [26] and RFPlasmid [27], and assembly-graph or homology-based tools such as plasmidSPAdes [28], SCAPP [29], Platon [30] and PlasForest [31]. These tools generally aim to replace or compete with the reconstruction step itself. PLAGUE, on the other hand, occupies a different group built on MOB-suite as a post-processing validation layer, so that MOB-suite’s typing, clustering and multi-contig binning, capabilities not all of the tools above replicate, are retained rather than discarded.

PLAGUE’s central principle is the integration of several established validation concepts - circularization detection, coverage estimation and discordant-read analysis - into a single, reproducible, post-processing framework. Rather than replacing existing plasmid reconstruction algorithms, PLAGUE leverages complementary evidence to evaluate the confidence of MOB-suite’s predictions while preserving their biological interpretation and per-plasmid organisation. A key design consideration was how coverage information should be used. PCN varies substantially across replicons, ranging from approximately one copy per chromosome for tightly regulated low-copy plasmids to tens or even hundreds of copies per cell for small multicopy elements. Moreover, PCN is not a static property; it can change dynamically in response to environmental conditions, host background, growth phase, and selective pressures [32]. Recent work has shown that transient plasmid amplification can increase plasmid copy number by one to two orders of magnitude, driving elevated gene dosage, enhanced horizontal transfer, and amplification-mediated heteroresistance. Consequently, coverage variation may represent biologically meaningful changes in plasmid abundance rather than assembly error [33]. For this reason, PLAGUE deliberately avoids applying hard coverage-based exclusion criteria to multi-contig plasmid predictions. Aggressive filtering based on coverage risks removing biologically relevant plasmids, particularly high-copy plasmids and amplified resistance plasmids that may be of greatest epidemiological interest. Instead, PLAGUE restricts coverage-based filtering to single-contig predictions, where coverage provides a more direct indicator of assembly support, while reporting estimated PCN as an informative annotation rather than a strict exclusion criterion.

The benchmark results confirm this substantial, quantifiable gains in precision at modest and predictable sensitivity cost. Aggregated across all four coverage tiers, PLAGUE reduced false positives in *E. coli* from 297 to 218, a 27% reduction, while retaining 98% of true positives (585 of 597). In *S. aureus*, the species with the highest artefact burden (unfiltered precision as low as 19.5-22.9%), PLAGUE removed 58% of false positives (421 to 176) while retaining 92% of true plasmids. *B. cereus*, PLAGUE reduced false positives by 32% (124 to 84) at an aggregated sensitivity of 87.0%, the lowest of the three species. Moreover, sensitivity fell further, to 78.9%, at the 10x stress-test depth, the one condition under which PLAGUE’s recall cost becomes non-trivial. Outside this worst case, PLAGUE’s aggregated sensitivity across species ranged from 87.0% to 98.0%, for false-positive reductions of 27-58%. Several limitations should nevertheless be acknowledged. For example, PLAGUE does not perform de novo plasmid assembly, improve the initial contig binning generated by MOB-suite, or overcome the intrinsic limitation of short-read plasmid reconstruction. Rather than replacing existing reconstruction methods, PLAGUE serves as a validation and refinement layer that increases confidence in reconstructed plasmids while preserving the rich annotation and per-plasmid structure required for downstream evolutionary and epidemiological analyses.

Benchmarking against geNomad [7], a sequence-based classifier, shows that neither tool is categorically superior. The better choice depends on species and on whether sensitivity or precision is the priority. In *E. coli*, PLAGUE outperformed geNomad on both sensitivity (98.0% vs 91.8%) and F1 (0.836 vs 0.823), at a marginally lower precision (72.9% vs 74.6%). In *S. aureus*, PLAGUE’s sensitivity advantage was much larger (92.0% vs 50.9%) but came at a substantial precision cost (36.9% vs 71.3%). The combination of both tools resolved this trade-off, reaching 91.1% sensitivity, 76.1% precision and F1 = 0.829, the best result obtained for this species by any configuration. In *B. cereus*, geNomad alone was the stronger tool on every axis, precision, sensitivity and F1 (0.772 vs 0.669), indicating that for this species sequence composition alone is more discriminating than PLAGUE’s structural evidence, and that combining the two tools did not recover this gap. PLAGUE and geNomad should therefore be read as complementary evidence sources. Hence, the combined mode is recommended specifically for species with high frequency of artefacts such as *S. aureus*, while geNomad alone, or PLAGUE with relaxed thresholds, is a reasonable alternative for species such as *B. cereus*.

This benchmark covered three species chosen to span Gram-negative and Gram-positive bacteria and a range of plasmid architectures and artefact prevalence, but it remains a three species validation, and generalisation should be stated carefully. The argument for broad applicability is strongest for Enterobacteriaceae, a family that shares a large, overlapping pool of circulating plasmid backbones [34], which is precisely the structural signal, replicon, relaxase and mobility conservation, that both MOB-suite and PLAGUE’s evidence layers rely on, so performance in *E. coli* is a reasonable proxy for related genera such as *Klebsiella* or *Salmonella*. That argument does not extend automatically to *S. aureus* and *B. cereus*, which are single-species representatives of Gram-positive plasmid biology tested here specifically because their filtering behaviour differed most from *E. coli* and performance in other Gram positive pathogens, or in taxa with markedly different plasmid architectures such as *Acinetobacter* or *Pseudomonas*, has not been tested. The filter thresholds themselves were derived empirically from this benchmark and from Illumina data specifically, and may require recalibration for other sequencing platforms, library preparations, or coverage distributions.

It is also important to acknowledge limitations in the reference ground truth itself. Benchmarking against NCBI RefSeq assumes that reference assemblies are complete and correctly annotated, which is not always the case for a class of elements as dynamic as plasmids [1] [2]. Some predictions scored as false positives here may therefore be genuine but unannotated extrachromosomal elements, transient amplifications, or resistance plasmids simply absent from the reference used, meaning the precision gains reported above are, if anything, a conservative estimate of PLAGUE’s true performance.

Beyond accuracy, PLAGUE is built for routine, at-scale use. The pipeline is implemented in Nextflow [10] and packaged as a single command that runs MOB-suite, read mapping and filtering as one workflow. Containerisation via Docker or Singularity keeps this behaviour consistent across systems, and the same pipeline definition runs unchanged on a local workstation, an HPC cluster under Slurm, or cloud infrastructure. Nextflow also parallelises independent samples automatically, which improves throughput when screening many isolates at once. This does not, however, change the per-sample computational cost, where PLAGUE’s peak memory use is roughly six-fold lower than geNomad’s, but its wall-clock runtime is up to four-fold longer, making it best suited to modest compute environments where memory, not time, is the limiting factor (Fig. 3). All intermediate files are retained by default so filtering decisions remain traceable, though users can discard them once a run is finalised.

Taken together, PLAGUE improves confidence in MOB-suite-derived plasmid reconstructions using transparent, auditable, data-supported metrics, at a precision gain of 27-58% in false-positive reduction and a sensitivity cost that remained at or below 13% in aggregate across all three species, rising to a maximum of 21% only at the 10x stress-test depth, the least favourable condition tested. This profile is best suited to the settings where short read sequencing continues to dominate such as, AMR surveillance, hospital and routine clinical genomics, retrospective mining of public repositories, and outbreak investigation, contexts in which both false positives and missed plasmids carry a real analytical and public-health cost.

## Conclusion

Short reads are still the vast majority of available sequencing data for bacteria, but it poses challenges to well-established pipelines such as MOB-suite. PLAGUE is a post-filtering optimized for decreasing the frequency of false positives under suboptimal coverage, while not discarding real plasmid predictions. PLAGUE performs comparably to geNomad, while showing less variation in regards to the bacterial species background and requiring less computing power.

## Key points

- PLAGUE is a post-processing validation layer for MOB-suite, not a replacement for it: across three benchmarked species it removed 27-58% of false-positive plasmid predictions while retaining 87-98% of true positives.
- Its conservative, multi-criterion filtering (circularisation, coverage, discordant-read evidence) requires two independent lines of evidence to fail a prediction, keeping sensitivity high even in heavily fragmented assemblies.
- Performance is robust from full sequencing depth down to a 10x stress test, with the largest absolute false-positive reductions occurring at the lowest, most error-prone coverage tiers.
- PLAGUE and the sequence classifier geNomad are complementary rather than competing. PLAGUE has the sensitivity advantage in *E. coli*, the combined mode performs best in *S. aureus*, and geNomad alone is preferable in *B. cereus*.
- PLAGUE has a substantially lighter peak memory footprint than geNomad (about 6-fold lower) at the cost of a longer runtime (up to 4-fold), making it well suited to standard servers and routine surveillance workflows without high-memory infrastructure.

## Data availability

Accession ID to all datasets used in this work are presented in supplementary Supplementary Tables S3-S5.

---

**Figure S1:**
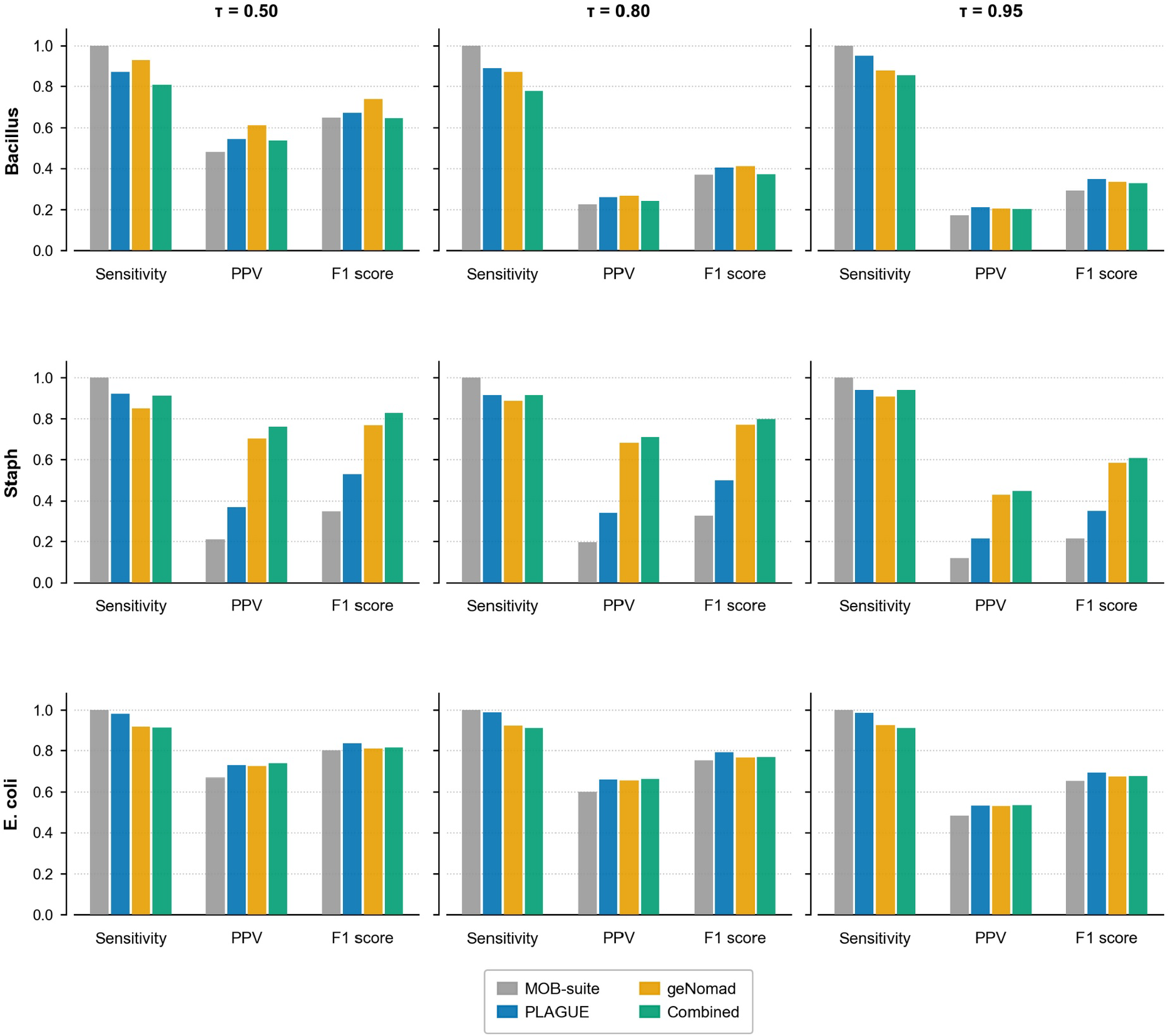
Sensitivity, positive predictive rate and F1 scores for different target species and cutoffs of containment (τ). Metrics are derived from containment to the reference (CR = |B∩R|/|R|) and containment on the assembly (CA = |B∩R|/|B|), where B is a MOB-suite bin and R is an NCBI reference plasmid. To derive a single summary value per MOB-suite bin, the ratio CR/CA = |B|/|R| was computed, capturing the relative size of the predicted bin with respect to the reference. The ratio thresholds for assigning a construction as a true or false finding were: tau = 0.50, 0.80, 0.95.

